# A phase field model with stochastic input simulates cellular gradient sensing, morphodynamics, and fidelity of haptotaxis

**DOI:** 10.64898/2026.03.10.710962

**Authors:** Joseph M. Koelbl, Jason M. Haugh

## Abstract

Haptotaxis is an understudied form of directed cell migration in which movements are biased by gradients of immobilized ligands. For example, fibroblasts and other mesenchymal cells sense and respond to gradients of extracellular matrix (ECM) composition, which is relevant during tissue morphogenesis and repair. As a step towards understanding how haptotactic gradients spatially bias cell adhesion, intracellular signal transduction, and cytoskeletal dynamics, we formulated a phase field model of whole-cell migration, in which the occupancy of potential adhesion sites changes stochastically with time. With careful assignment of parameter values, the model predicts significant haptotactic bias for adhesion-site gradient steepness of a few percent across the cell. We then used the model to predict how the cell’s removal of surface-bound ECM ligand (as observed in experiment) and/or the presence of a competing, chemotactic gradient influence(s) haptotactic fidelity. An emergent principle is that gains in directional persistence naturally offset losses of directional bias, at the cost of greater cell-to-cell heterogeneity of the response. In the case of orthogonally oriented gradients, this offset manifests as a remarkable robustness of the multi-cue response.

## INTRODUCTION

Directed cell migration guided by chemical and other spatial cues is required for embryonic development, tissue regeneration and homeostasis, and immune responses in vertebrates (1–6). In this field, migration biased by soluble compounds (chemotaxis) and the associated signal transduction have been broadly studied; fast-moving, amoeboid cells such as neutrophils (4–10) predominantly migrate by chemotaxis. In contrast, mesenchymal cells such as fibroblasts migrate much slower and can be directed by various types of spatial cues. These include chemotactic and haptotactic gradients, the latter defined as gradients of immobilized ligands, as found in the extracellular matrix (ECM) (11, 12). Both the relatively slow migration and haptotactic sensing of mesenchymal cells derives from their persistent adhesion to ECM, which is their natural microenvironment; the specificities of ECM adhesion are controlled via differential expression of cognate adhesion receptors of the integrin family (13, 14).

Relative to chemotaxis, haptotactic signaling and migration are not yet well understood. Despite foundational papers that have implicated certain molecular pathways associated with fibroblast haptotaxis (15, 16) and adhesion-based signaling therein (17), we lack mechanistic understanding of the relationships between subcellular processes and cell motility/shape changes (morphodynamics). In the general context of adhesion-based migration, we know that clusters of ECM-bound integrins form nascent adhesion structures, which physically engage the actin cytoskeleton while also mediating downstream signaling that modulates actin polymerization, turnover, and contraction dynamics (14, 18, 19). Thus, it would seem that haptotactic sensing and the mechanics of the haptotactic response are coupled. Attempts to integrate these dynamics at different scales of complexity have been approached through modeling. Published models of chemotactic signaling (20, 21) and integrin-mediated signaling (22, 23) in mesenchymal cells have been offered, as have relevant models of single (24) or collective (25) cell motion. A key missing link is the general relationship between adhesion-based signaling and cell morphodynamics, much less how that relationship is altered in the context of haptotaxis. In that regard, an essential aspect to consider is the apparently stochastic nature of discrete nascent adhesion formation, disintegration, and maturation events, which have been empirically and theoretically associated with local, leading-edge signaling and motility dynamics, without regard to haptotaxis (22, 26, 27). To bridge these, we show that stochasticity of adhesion formation in proximity to the leading edge, combined with a deterministic, phase field model of adhesion-based sensing and motility, predicts whole-cell haptotactic gradient sensing, attendant morphodynamics, and directional haptotactic responses.

Mathematical modeling of whole-cell migration, in a manner that integrates subcellular/local dynamics and whole-cell behavior, is not trivial. In addition to the aforementioned complexities, one must also confront how to handle updating the cell membrane/boundary based on predicted local stresses and assumed details of the cell mechanics. Some methodologies to probabilistically update the boundary of the migrating ‘cell’ include cellular Potts modeling (28, 29) and voxel-based finite element modeling (30). Other models update the cell edge on a schedule that is based on an actin force-velocity relationship (31). A distinct approach is to remove the explicit definition of the cell boundary, using the *phase field* formalism, which effectively replaces a conformal boundary of the cell with the gradient of an auxiliary variable that delineates the cell ‘phase’ from rest of the simulated domain. Originally developed to describe multiphase systems in materials science (32–34), the method has found application in the biological sciences, including in studies of vesicle dynamics (35, 36), symmetry breaking in cell migration (37, 38), keratocyte adhesion (39), membrane tension (40), and *D. discoideum* chemotaxis (41).

Here, we incorporate the phase field approach to develop the first model of haptotaxis that duly predicts signaling dynamics, cell morphodynamics, and cell migration behavior. An inherently distinct and necessary aspect of the model is stochastic implementation of the haptotactic gradient input, which probabilistically assigns (and later unassigns) nascent adhesion sites at each time step. This simple formulation recognizes the discrete and practically immobile nature of adhesion structures. The submodel of stochastic nascent adhesion dynamics feeds into a deterministic, phase-field model of the cell response, with simplified representations of adhesion-based signaling and regulation of actin-based protrusion. With appropriate parameterization of the model, simulations predict haptotactic fidelity, with adhesion gradients of >5% difference across a typical cell diameter yielding reliable haptotaxis. We next applied the model to generate testable predictions along two, distinct lines. First, we used the model to predict how removal of the ECM ligand from the surface (42) influences haptotactic behavior; and second, we explored the influence of a chemotactic cue opposing or oriented orthogonal to the haptotactic gradient. In all cases, an emergent principle is that gains in directional persistence naturally offset losses of directional bias (relative to the initial, haptotactic gradient), at the cost of increased cell-to-cell heterogeneity of the response. In the case of the orthogonal gradients, this offset manifests as a remarkable robustness of the multi-cue response.

## RESULTS

### A new model of mesenchymal haptotaxis on ECM with stochastic adhesion dynamics

Our haptotaxis model couples two distinct submodels (**Figure 1**). The first is a coarse-grain stochastic model that designates sites of potential nascent adhesion formation on a planar grid (**Fig. 1A&B**). We varied the probability of adhesion formation at the initial cell centroid position, expressed as a percentage. The gradient of the probability is exponential in the vertical direction, such that the % change across a distance equal to the initial cell radius (the arbitrary unit length of the simulation) is constant; different values of the % gradient, together with the no gradient/uniform probability, were assessed. At each time step, a certain number of nascent adhesion sites are assigned based on the probability, with turnover of the sites based on a mean lifetime of 60 s, a value consistent with experimental measurements (43). The stochastic output of this submodel, which defines the nascent adhesion variable *n*(*x,y,t*), is saved for potential reuse and comparisons among different variations of the second submodel.

**Figure 1:**
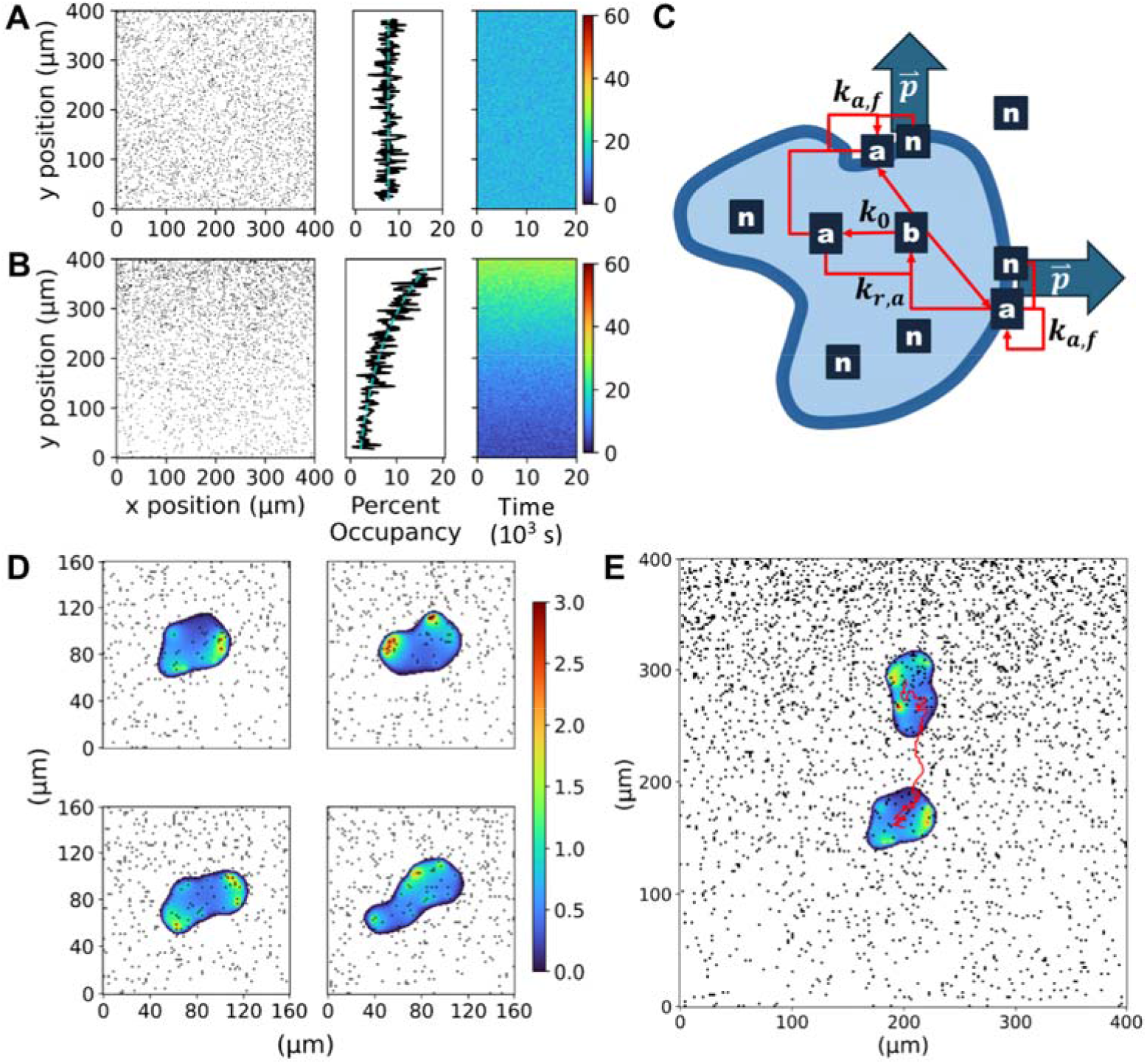
Phase field model of haptotactic sensing and migration. **A&B)** Snapshots of stochastic nascent adhesion occupancy (black on white) are shown for: 7.5% adhesion occupancy with no gradient (A) and 5% initial adhesion occupancy with 10% relative gradient (B). Side-bar plots show the fraction of sites occupied in each *y*-position row, compared to the generating exponential curve, and a kymograph of the same. **C)** A schematic of the phase field model depicts potential nascent adhesion sites, *n*, that can appear throughout the modeling space. Adhesions at the periphery mediate localization of the activator signaling protein, *a*, causing local, outward advection of the phase field through the protrusion vector, **p**. **D&E)** For one of the simulations corresponding to panel B, a sample of cell snapshots are shown in the 8×8 modeling inset, with activator value *a* indicated by the color scale (D); early and final cell states, and the 4.3-hour centroid track, are shown overlaid on the final adhesion occupancy (E).

The second submodel is a set of partial differential equations, adopting the phase field modeling formalism, which determines the cell morphodynamics and migration as a response (shape changes and translocation of the centroid) to the dynamics of the nascent-adhesion input. We found that the finite volume method (implemented in the python package FiPy) was suitable for the hybrid stochastic/deterministic model explored here. **Fig. 1C** illustrates the phase field, represented by the light-blue intracellular region (*ϕ* = 1) and dark-blue transition region (∇*ϕ*), and its resident variables. The submodel assumes that nascent adhesions (*n*), while in the transition region at the cell periphery, mediate the autocatalytic generation of activator (*a*), a proxy for adhesion-based signaling pathways such as GTP-loading of the Rho-family GTPase Rac1 (activated by guanine-nucleotide exchange factors like Tiam1 (44, 45) and β-Pix (46)) and phosphoinositide 3-kinase (PI3K)-dependent generation of the 3’ phosphoinositide, PIP_3_ (17), both of which promote Arp2/3-based dendritic actin assembly through WAVE, which has been shown to be essential for haptotaxis (15, 16, 47, 48). The activator species is formed from an inactive/precursor state (*b*); its concentration is considered spatially uniform/well-mixed and set by overall mass conservation. The role of the activator is to mediate, along with nascent adhesions, the outward protrusion of the transition region (velocity vector **p**), which results in deformation/shape changes of the cell and overall cell translocation in each simulation (**Fig. 1D&E** and **Video 1**). This vector variable represents actin protrusive force on the membrane, as previously considered (40). Our formulation, similar to previous models (23, 26) recognizes that actin-based protrusion relies on both actin polymerization (governed by *a*) and mechanical clutching (governed by *n*). The ‘diffusion’ of the associated force is analogous to viscous dissipation but also reflects the spread of F-actin barbed ends via Arp2/3-mediated branching (15, 23).

Qualitatively at least, the model structure and its parameterization produce simulations that are consistent with observed dynamics in migrating fibroblasts (**Figure 2** and **Video 2**). During migration on a surface coated with ECM ligand, such as Fibronectin (FN), clustered adhesion receptors (integrins) mediate signaling and actin protrusion (17). These discrete adhesion structures were visualized in live fibroblasts expressing the marker, EGFP-Paxillin, by total internal reflection fluorescence (TIRF) microscopy, together with 3’ phosphoinositide signaling as monitored by recruitment of the biosensor, mScarlet-AktPH, to the plasma membrane; over a span of 4 minutes, there is modest translocation of the cell, but localized motility correlated with adhesion-based signaling dynamics can be substantial (**Fig. 2A**). A representative montage from simulation results, spanning the same time period, shows similar dynamics (**Fig. 2B**).

**Figure 2:**
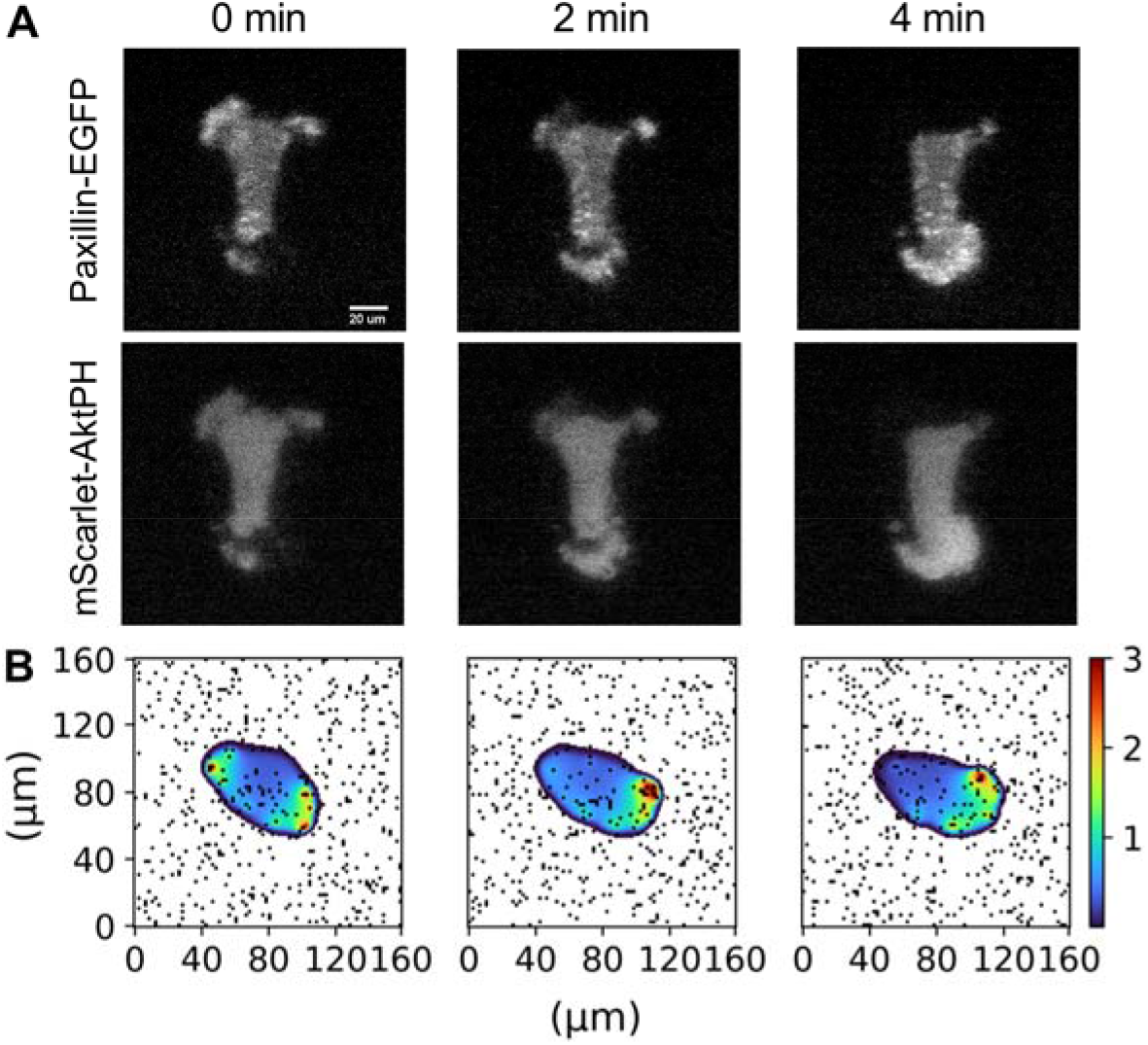
Dynamics of adhesion-based signaling. **A)** Montage of experimental TIRF images of a migrating fibroblast taken over a brief sequence spanning 4 minutes. Two-color imaging of Paxillin-EGFP, marking adhesion structures, and of mScarlet-AktPH, a translocation biosensor of PI3K signaling. The sequence shows local dynamics of adhesion, signaling, and shape change on this time scale. **B)** Over the same time span, the simulations show qualitatively similar dynamics.

### Simulated haptotaxis is sensitive to ECM density and gradient steepness

To assess the haptotactic fidelity of the simulated cells, we varied the simulated ECM gradient properties (initial adhesion density and the relative steepness) and quantified the cells’ directed migration responses in terms of the Forward Migration Index (FMI) (**Figure 3**). According to this broadly used metric, a value of 1 corresponds to perfectly persistent attraction towards the gradient, whereas a value of −1 corresponds to perfect repulsion. As mentioned above, we varied the probability of adhesion formation at the initial cell centroid position, with three values (2.5%, 5%, and 7.5%); and we varied the relative gradient steepness, with three values (2%, 5%, and 10%) together with no gradient/uniform probability. For each of those 12 gradient conditions, a total of 18 simulations were performed, with 6 different *n*(*x,y,t*) schedules (different supersets of random numbers) and 3 different *x*_0_ starting positions each.

**Figure 3:**
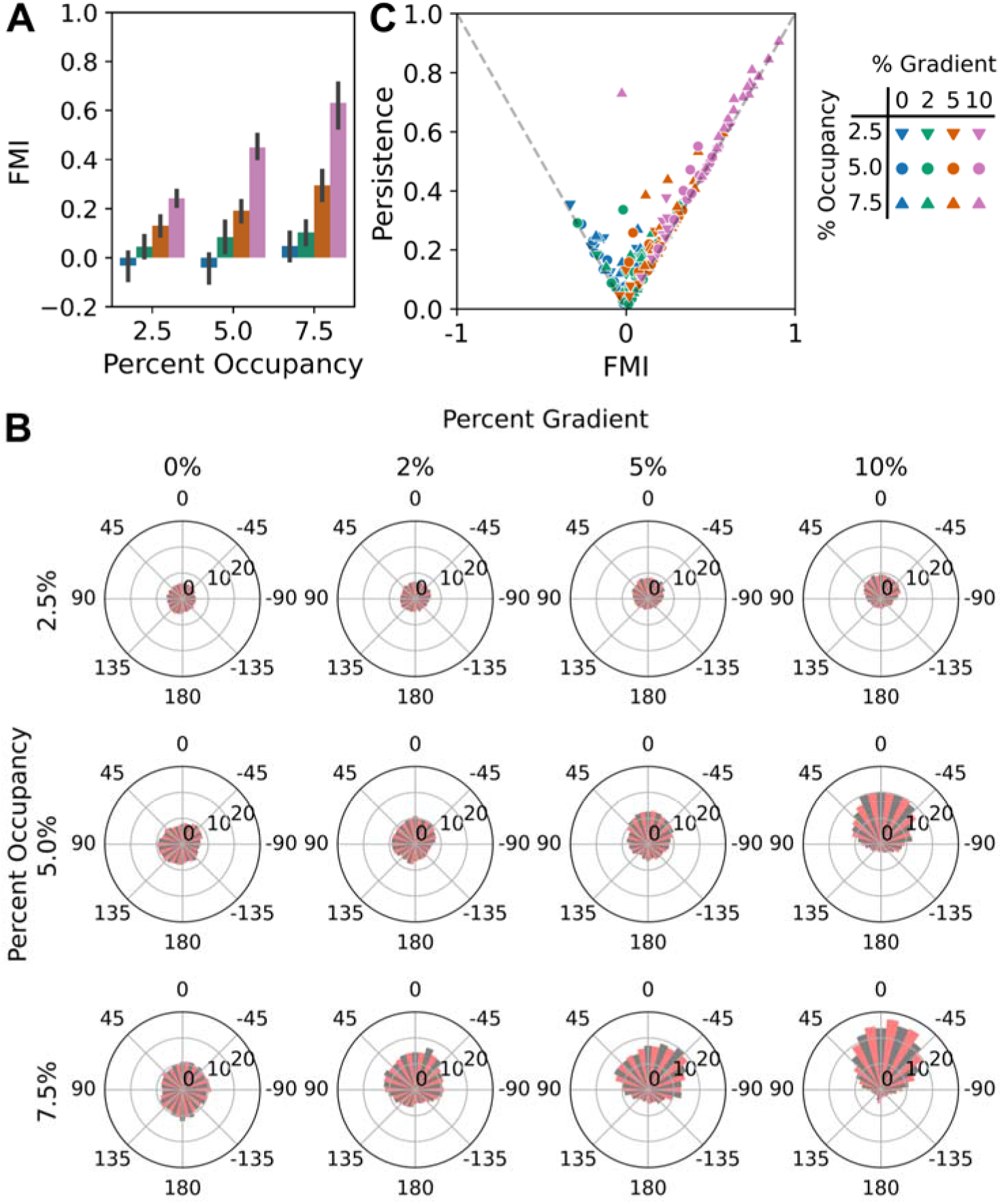
Analysis of haptotactic fidelity predictions. **A)** The haptotactic fidelities of modeled cell-centroid tracks were quantified in terms of the standard Forward Migration Index (FMI), varying the random number generation and cell starting position (N = 18) for each gradient condition (initial adhesion occupancy and relative gradient as indicated). FMI values are plotted as mean ± 95% confidence interval. **B)** For each gradient condition, wind rose plots show the polar histograms of cumulative centroid displacement, as a function of centroid movement angle relative to the gradient. **C)** For each simulation, symbolized by gradient condition as shown, the FMI value is decomposed as the product of the persistence value (straightness of the migration track) on the vertical axis and the cosine of the displacement angle relative to the gradient.

The results show that gains in the mean FMI reflect increases of both the initial adhesion density and the relative gradient (**Fig. 3A**). For the model parameterized as it was, haptotaxis was generally significant for relative gradient values of 5%, but also for 2% relative steepness when initial gradient density was sufficiently high. As explained under Methods, one may decompose the value of FMI as the product of the Persistence ratio (commonly referred to as D/T, a metric of migration path ‘straightness’) and the sine of the displacement angle *θ*, which indicates where the cell ended up relative to the gradient direction. A plot of Persistence vs. FMI indicates the relative contributions of those effects (**Fig. 3B**), with higher adhesion density promoting greater persistence and higher steepness promoting better final alignment with the gradient. Another common way to display directed migration data is as a Wind Rose plot, a polar plot of cell-centroid displacements as a function of angle relative to the gradient. The plots for the various gradient conditions illustrate the intensification of the migration towards the gradient as both initial adhesion density and gradient steepness are increased (**Fig. 3C**).

### ECM removal by cells enhances haptotactic fidelity via increased persistence

Having established the sensitivity to ECM gradient conditions in our ‘base-case’ model, we turned to the question of how dynamic changes to the ECM gradient field, brought about by the cells’ removal of ECM ligands from the surface, might influence the haptotactic response (**Figure 4**). (49)Evidence of FN removal by fibroblasts has been long documented for uniform FN coating conditions (49–54). As explained under Methods, we considered that the presence of the cell (phase field) occupying the surface permanently ‘degrades’ the probability of adhesion-site formation at underlying locations. An example of a simulation with ECM removal is shown in **Fig. 4A&B** and **Video 3**. The snapshot shown in **Fig. 4B** shows the positions of the adhesion sites, which by inspection have a much lower density within the denuded region compared with that of the surrounding area.

**Figure 4:**
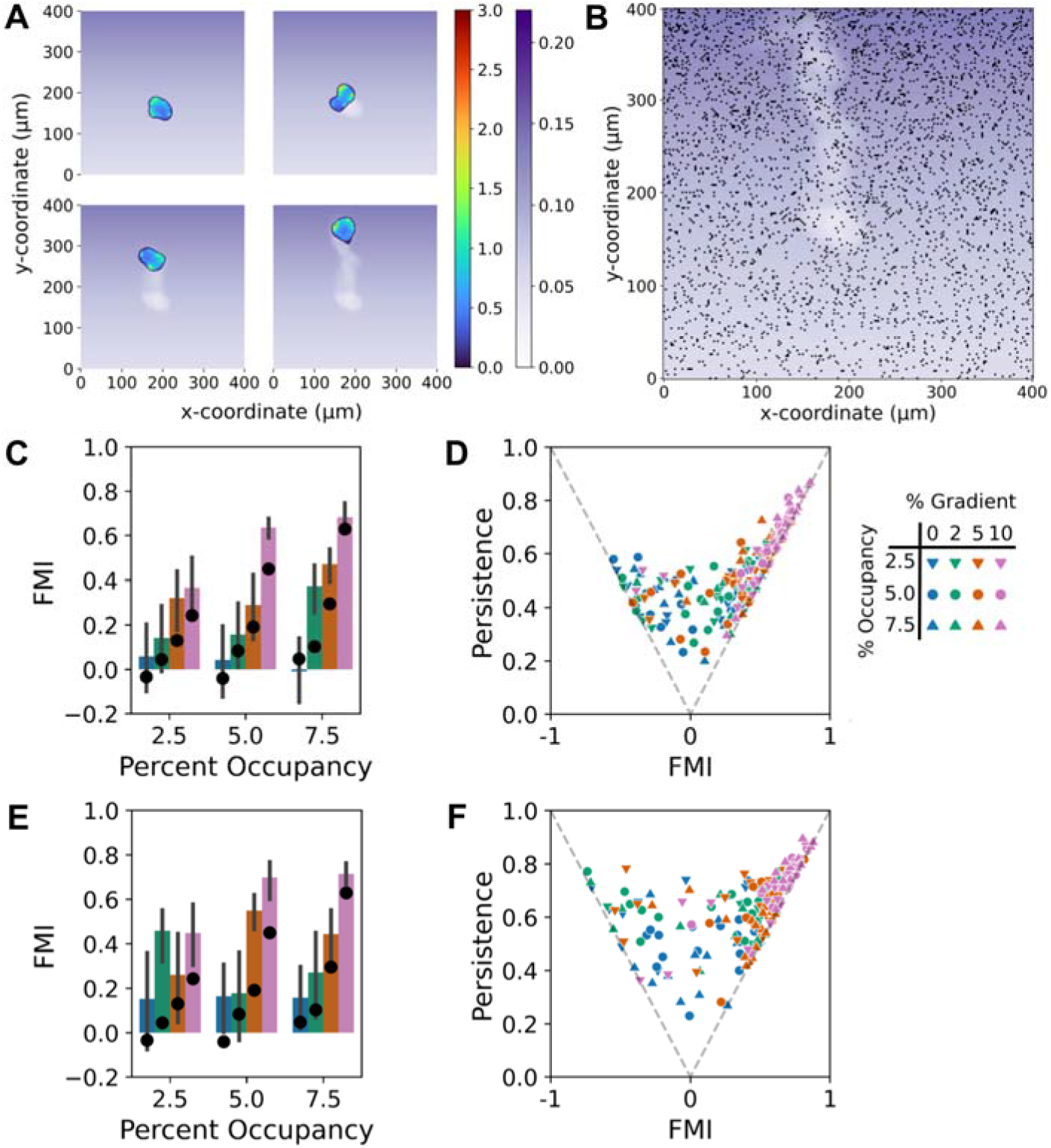
ECM removal by cells enhances haptotactic fidelity via increased persistence. **A)** Montage showing how the probability of adhesion site selection (grayscale) is degraded according to occupancy by the phase field cell (activator value shown in color scale). **B)** The same cell track is replotted without the cell and overlaid with the adhesion site occupancy at the final time point, showing reduced density of adhesion sites in the cell path. **C&D)** With low removal rate, FMI and persistence results are plotted as in Fig. 3A&C (N = 18 for each gradient condition). In C, the mean FMI values without removal, from Fig. 3A (black circles), are shown for comparison. **E&F)** Same as C&D, but with high removal rate.

In principle, ECM removal allows cells to either steepen the haptotactic gradient or generate their own, competing ECM gradient. To assess the effect on haptotactic fidelity, simulations with low and high removal rates were carried out under the same adhesion-site gradient conditions explored above. The results show that, relative to no removal, adding a low removal rate tends to increase the mean but also the variability of the FMI, calculated based on the initial gradient direction (**Fig. 4C**); clear improvements were the ability to sense a very shallow (2%) gradient at the highest initial adhesion density and more robust response to steeper (≥ 5%) gradients. Further analysis reveals that the predominant effect of the lower removal is an increase in cell Persistence ratio (**Fig. 4D**; compare with **Fig. 3B**), which amplifies both the mean and variability of the FMI; under zero or shallow gradient conditions, some simulations exhibited substantially negative FMI values. The same qualitative conclusions apply for the higher removal rate (**Fig. 4E&F**); cell persistence is further increased, yielding more extreme and less predictable haptotactic outcomes.

### Opposing, anti-parallel haptotactic and chemotactic gradients reveal their relative strengths of attraction

During wound healing, fibroblasts are exposed to both chemotactic and haptotactic cues (11); therefore, a natural extension of the model was to assess dual gradient sensing. While it is intuitive that co-aligned chemo- and hapto-tactic gradients ought to enhance cell migration toward the gradients’ direction, and that anti-parallel gradients ought to frustrate directed migration, our model may be used to quantitatively assess such effects. To address this question, we modified the model to allow for a chemotactic receptor to mediate generation of the activator species, in parallel with *n*. Unlike the adhesion sites, which arise stochastically, activation of the chemotactic receptor follows a deterministic function of its gradient direction.

With this modification in place, we first evaluated opposing, anti-parallel gradients to most directly compare gradient potency (**Figure 5**). We tested combinations of zero, medium (5%), and high (10%) gradient steepness for each gradient type, for the usual initial adhesion density values (2.5%, 5%, 7.5%), and we also varied the degree of ECM removal (none vs. low). Each of the 54 new conditions was replicated for 12 random number supersets, and the results are shown alongside conditions with no chemotactic signaling (as shown in Figs. 3 & 4). For each simulation, the FMI was calculated for the haptotactic direction (Haptotactic Index, or HI), with negative values indicating dominance of chemotactic sensing. As illustrated for gradients with 5% steepness in opposite directions, we found combinations of gradient conditions that frustrated cell migration (**Fig. 5A** and **Video 4**); indeed, for base-case adhesion gradient conditions that otherwise predict significant haptotaxis, a 5% chemotactic gradient effectively neutralized tactic migration, whereas a 10% chemotactic gradient was dominant (**Fig. 5B**). These results quantify how the opposing gradients thumb the scale towards a haptotactic, neutral, or chemotactic outcome. With low ECM removal, the conclusion is largely the same, with the notable qualitative differences at 2.5% adhesion occupancy (**Fig. 5C**); under this condition, chemotactic signaling boosts the overall magnitude of the otherwise relatively weak activator generation, actually promoting haptotaxis when the relative steepness of the haptotactic gradient exceeds that of the chemotactic gradient (compare **Fig. 5B&C**). Besides these qualitative differences, the quantitative effects are as described in the previous section: amplification of mean and variance of tactic fidelity, through increased migration Persistence ratio (**Fig. 5D**). Here, the amplification of fidelity applies to either haptotactic or chemotactic dominance.

**Figure 5:**
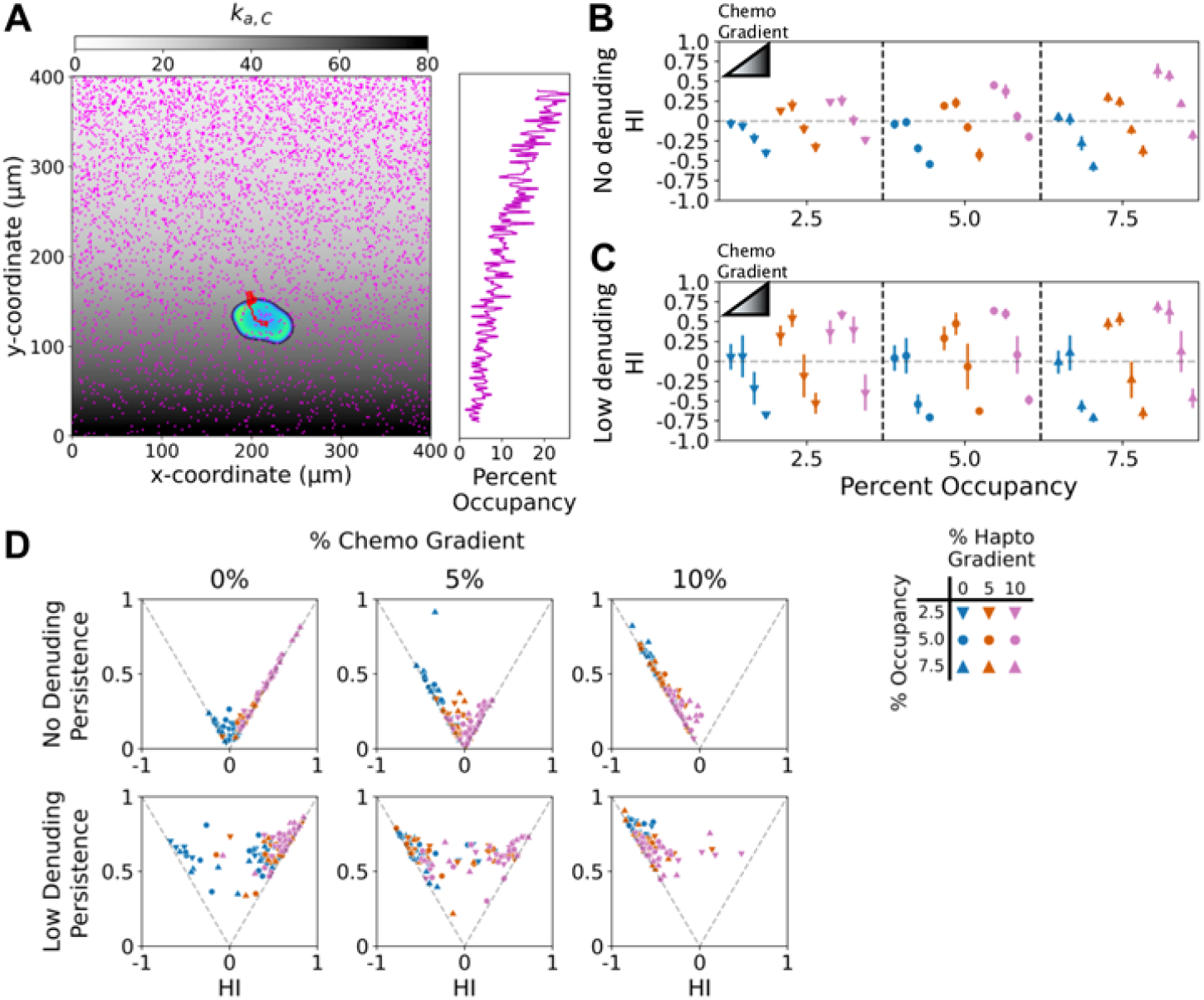
Frustration of haptotaxis by an opposing chemotactic gradient. **A)** A simulated cell responding to opposing haptotactic and chemotactic inputs, each with 10% relative gradient, is displayed with its cell track. The final adhesion occupancy is shown in magenta, and the magnitude of the chemotactic input, *k*_*a,C*_, is overlaid in grayscale; the side bar plots the fraction of sites occupied in each *y*-position row. **B&C)** To distinguish between opposing haptotactic and chemotactic directions, the FMI in the haptotactic direction is called the Haptotactic Index (HI). HI values are reported as mean ± 95% confidence interval (N = 12) for each combination of haptotactic (initial adhesion occupancy and relative gradient as shown) and chemotactic (none or chemoattractant with 0, 5, or 10% relative gradient) conditions. Simulations were run with no ECM removal (B) and low removal (C). **D)** Contribution of cell persistence to the HI values.

### Cells exhibit robust, bidirectional fidelity to orthogonal chemotactic and haptotactic gradients

Having established and characterized a simple dual-gradient model, we used it to predict the response to orthogonally oriented hapto- and chemo-tactic gradients as a uniquely insightful experiment (**Figure 6**). We simulated the same set of conditions as described in the previous section, but with the chemotactic gradients oriented in the *x*-direction. With each gradient oriented on its own axis, the FMI values for the two perpendicular directions may be calculated; here we refer to these as the aforementioned HI and the Chemotactic Index (CI). As shown in **Fig. 6A** and **Video 5**, with steep gradients in both directions, cells are able to sense and migrate towards both spatial cues. But, to what extent does the presence of a chemotactic gradient ‘distract’ the cell from adequately sensing the haptotactic gradient, and vice versa? Intuitively, one might expect that chemotactic conditions that neutralize haptotaxis or reverse tactic dominance when presented in the opposite directions would significantly, if not dramatically, alter haptotaxis when presented orthogonally.

**Figure 6:**
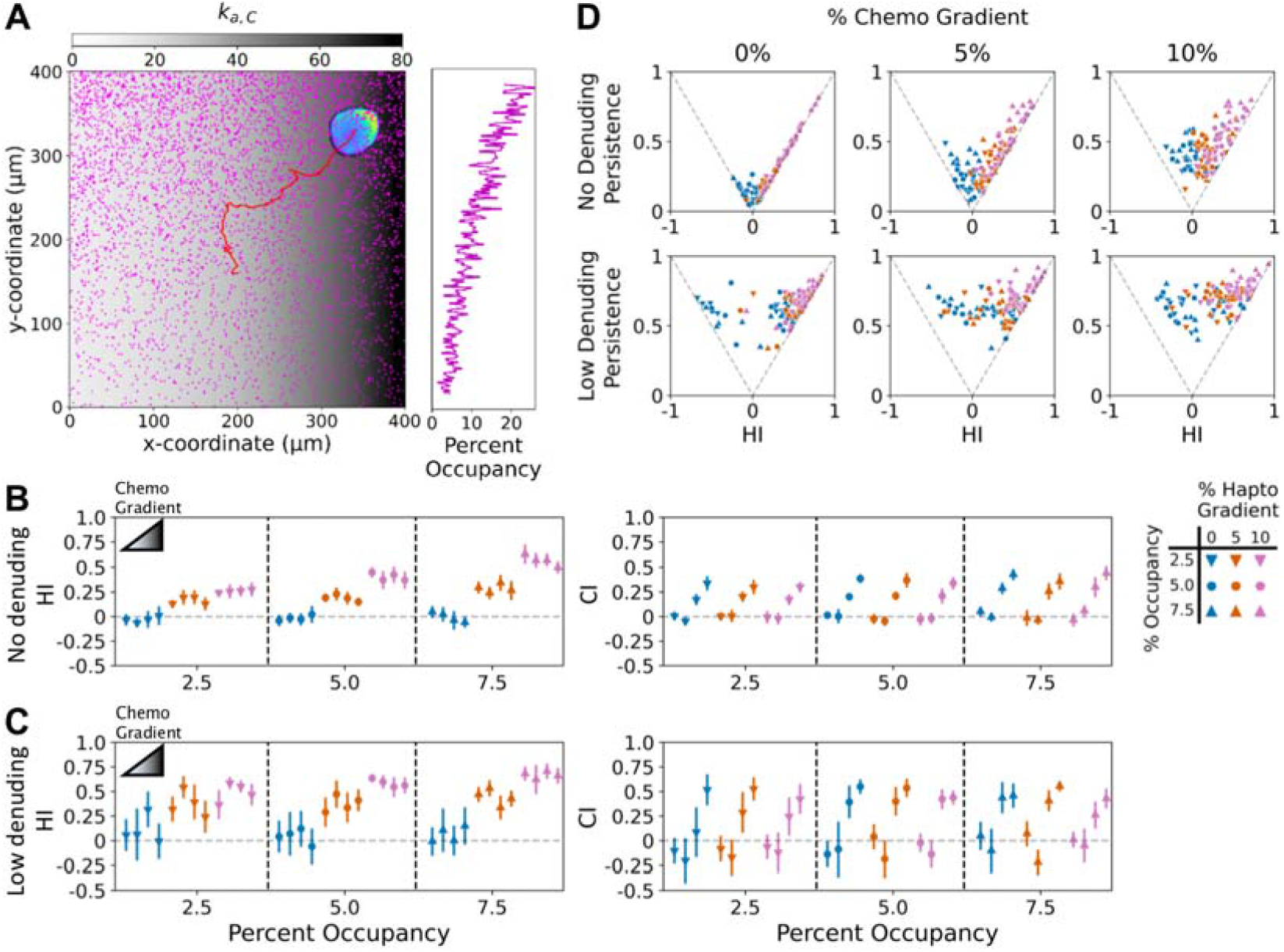
Robustness of the multi-cue response to orthogonal haptotactic and chemotactic gradients. **A)** A simulated cell responding to orthogonal haptotactic and chemotactic inputs, each with 10% relative gradient, is displayed with its cell track. The final adhesion occupancy is shown in magenta, and the magnitude of the chemotactic input, *k*_*a,C*_, is overlaid in grayscale; the side bar plots the fraction of sites occupied in each *y*-position row. **B&C)** To distinguish between orthogonal haptotactic and chemotactic directions, *y* and *x*, the FMI in the *y*-direction is called the Haptotactic Index (HI), and, in the *x*-direction, the chemotactic index (CI). These values are reported as mean ± 95% confidence interval (N = 12) for each combination of haptotactic (initial adhesion occupancy and relative gradient as shown) and chemotactic (none or chemoattractant with 0, 5, or 10% relative gradient) conditions. Simulations were run with no ECM removal (B) and low removal (C). **D)** Contribution of cell persistence to the HI values.

Yet remarkably, in the absence of ECM removal, the orthogonal, chemotactic gradient did not significantly influence haptotactic fidelity as measured by HI; and by the same token, the orthogonal, haptotactic gradient did not significantly influence the CI, even when the competing gradient was steeper (**Fig. 6B**). It is as if the two sensing systems were independent. With low ECM removal, the conclusion is almost the same (**Fig. 6C**), although the addition of uniform chemoattractant (with zero gradient) boosted HI in some cases (as seen also in **Fig. 5C**); here again, there was no significant decrease in HI or CI imposed by the presence of the other gradient.

The results presented above suggest that the overall enhancement of activator generation mediated by the two receptor types compensates for the competing orientations of the gradients. Consistent with this notion, analysis of Persistence ratio vs. HI shows that the mean HI is approximately maintained, despite deviation of the cells’ directionality by the chemotactic gradient, by increasing the persistence of migration (**Fig. 6D&E**).

## DISCUSSION

The model presented here is a new approach to assess hypotheses about haptotactic gradient sensing and resulting migration behavior, which is relevant for biased migration of fibroblasts and other mesenchymal cells on ECM with gradients of adhesive ligands. The stochastic input to the phase field model reflects the stationary nature of adhesion structures, in stark contrast to the ‘noisy’ input more appropriately assumed in models of stochastic chemotactic sensing (55, 56). A similar stochastic adhesion formalism has been employed in other models, with adhesions assigned to locations at random, and the adhesions acting as mechanical springs linking the intracellular cytoskeleton and ECM (39, 41); in those contexts however, the adhesions are not linked to intracellular, pro-protrusion signaling. Our simulations as parameterized predict the capability of sensing shallow haptotactic gradients as low as 2% steepness and robust haptotaxis at 5% steepness for the range of adhesion densities tested. Under these shallow gradient conditions, the associated FMI values align well with previously reported values for fibroblasts (15, 16); with higher steepness (10%), the simulations consistently ‘outperform’ observed haptotaxis.

Another novel aspect explored here is the known ability of fibroblasts to remove adhesive ligands from the surface as they migrate (49–54). Implementing the simplest model of this, based on the dwell time of the cell phase, the general trend is an increase in cells’ persistent directionality of migration. Insofar as a cell explores and alters the gradient direction, differing from the prescribed one, this results in a spurious trajectory that contributes to an increase in variability of the FMI; yet, under favorable conditions, more trajectories achieve deeper advances toward the gradient, consistent with experiment (15, 16).

Another, perhaps surprising prediction of the model concerned the response to two spatial cues, with a deterministic, chemotactic gradient presented either opposite or orthogonally to the stochastic, haptotactic one. Noteworthy in that context is that physical engagement of the protrusion ‘machinery’ still implicitly depends on the presence of nascent adhesions in the model. We also assumed that both gradient sensing inputs contribute to the local generation of the same activator variable. Earlier, we mentioned that adhesion complexes mediate Arp2/3-based dendritic actin assembly through activation of the PI3K/Rac/WAVE signaling axis as an essential haptotaxis pathway. This pathway is also activated in fibroblasts in response to the chemoattractant, PDGF (47, 57–61), though it is not strictly required for fibroblast chemotaxis (15, 62). Given this complexity, one may hypothesize that the two, co-presented cues could either synergize or interfere with each other. The present model predicts a surprising robustness of haptotactic fidelity when cells are challenged with an orthogonal chemotactic gradient, even when the strength of the competing gradient was judged to be greater.

Depending on one’s perspective, the simplicity of the present model might be viewed as either a strength or a major limitation. As we feel we have demonstrated, a simple model allows for analysis of basic yet novel concepts and may be better suited for generating qualitative predictions. On the other hand, from the perspectives of both signal transduction and cell mechanics, the model is limited by its simplicity and could be expanded. With an eye towards such an effort, we can point to phase field models of random motility that incorporate force dynamics on the membrane (35–39, 63) and to adhesion-based signaling models with similar force dynamics (22, 23). Additionally, we have already noted that there has been much work in the field focused on elucidating the detailed mechanisms of haptotactic and chemotactic signaling, which could be explored further. Of course, as one considers modeling what we know about signaling and cytoskeletal dynamics, one must also grapple with the need to formulate and specify associated model parameters. Per se, the exercise of refining a simple model is humbling and naturally reveals that which is required. We contend that it is a worthwhile endeavor.

## MATERIALS AND METHODS

### Stochastic simulation of nascent adhesion dynamics

Nascent adhesions (variable *n*) at a cell’s periphery activate the cell’s migration response. As the coarsest stochastic formalism, we place discrete sites on a planar, *N*_*x*_ × *N*_*y*_ grid and consider each site either an adhesion site ‘occupied’ (*n* = 1) or not (*n* = 0) at each time *t*. For each site, the fractional occupancy *χ*_*n*_ (expected value of *n*) is prescribed by a gradient in the *y*-direction, with constant relative steepness.

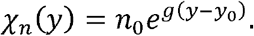

The two parameters introduced here are *n*_0_, the expected value of *n* at the cell’s initial centroid position (with *y* = *y*_0_), and g, the relative gradient steepness; both are expressed as percentages in the figures. Integrating *χ*_*n*_(*y*) over [0,*N*_*y*_*ℓ*], where *ℓ* is the lattice spacing, we obtain the expected number of occupied sites, *N*_*n*_.

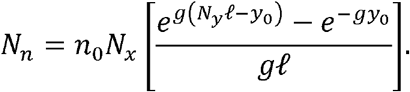

For random migration conditions, *g* = 0, and the limit value *N*_*n*_ = *n*_0_ *N*_*x*_ *N*_*y*_ applies.

To generate the site occupancy pattern for each time *t*, all sites are assigned a ranking score *s*_*n*_, according to its prescribed fractional occupancy and a random number rand drawn from [0,1).

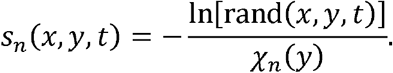

The initial values *s*_*n*_ (*x,y*,0) are ranked from lowest to highest, and sites with rank less than *N*_*n*_ + 1 are designated as occupied initially. During each time step, duration Δ*t*, a certain number of previously occupied sites are relegated to unoccupied status, replaced by the same number of previously unoccupied sites. That number is rounded from (Δ*t* / *τ*)*N*_*n*_, where the constant parameter *τ* is the mean lifetime of the adhesion site. The previously occupied sites that are switched to unoccupied are chosen at random, shuffled using “numpy.random.permutation”. These are replaced by previously unoccupied sites according to the lowest, updated *s*_*n*_(*x,y,t*).

For the simulations presented here, *τ* = 30 Δ*t*, with Δ*t* ~ 2 s in real time, and they were carried out for 10,000 time steps. According to this algorithm, each gradient *χ*_*n*_(*x,y*) and superset of random numbers yields a nascent adhesion input *n*(*x,y,t*), which was saved for each instance.

### Phase field model of adhesion-based sensing

The stochastic variable *n*(*x,y,t*) described above, is the input to a deterministic model of the cell response. In this model, the phase field equation explicitly models the interface between intracellular and extracellular ‘phases’ in planar geometry, with conservation of the auxiliary variable *ϕ* expressed as follows.

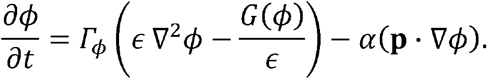

In the first term on the right-hand side, *Γ*_*ϕ*_ is a Lagrange multiplier, *ϵ* controls the thickness of the transition region between the phases represented by *ϕ*= 1 (intracellular space) and *ϕ* = 0 (extracellular space); phase separation is enforced by a double-well potential function, *G*(*ϕ*). The last term describes deformation of the transition region, marked by the gradient ∇*ϕ*, according to a protrusion velocity vector **p** and with *α* as a constant scaling parameter (40). The form of *G*(*ϕ*) used here is as follows.

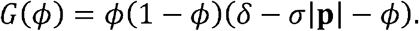

The function *δ* promotes area conservation relative to the initial phase field, *ϕ*_0_, with weight *μ*.

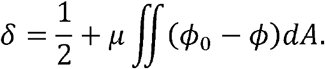

As shown in the expression for *G*(*ϕ*) above, the area conservation can be locally disrupted by the magnitude of **p**, with weight *σ* (40).

The protrusion velocity vector **p** is governed by the following force/momentum balance.

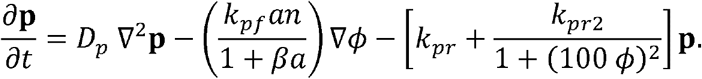

The first term on the right-hand side describes dissipation as the dendritic actin network spreads with constant ‘diffusivity’ *D*_*p*_ (analogous to kinematic viscosity). The second term describes the generation of outward protrusion force, directed opposite the phase-field gradient, dependent on peripheral signaling (*a*; see below) and mechanical clutching by nascent adhesion sites (*n*); it is saturable with respect to *a*, according to the constant parameter, *β*. The last, collective term describes the drag-like opposition of the protrusive force, with rate constants *k*_*pr*_ and *k*_*pr*2_; the latter contribution rapidly eliminates protrusion outside of the cell domain.

The activator of membrane protrusion is represented by concentration variable *a*, governed by the following conservation equation.

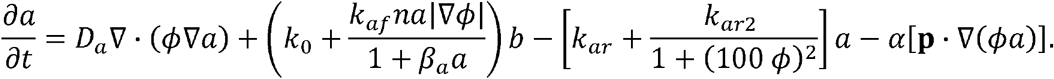

The first term on the right-hand side describes diffusion of the activator inside the cell phase, with constant diffusivity *D*_*a*_. The second, collective term describes generation of the activator, with conversion from an inactive form or precursor (concentration *b*), and is further explained below. The third, collective term describes reversion to the inactive/precursor state, with rate constants *k*_*ar*_ and *k*_*ar*2_, analogous to the **p** balance. The last term describes advection of the activator, consistent with the phase field balance.

The generation term in the activator balance above reflects both basal (*k*_0_) and autocatalytic signaling at the cell periphery (dependence on |∇ *ϕ*|), depending on cooperation of the activator (*a*) and adhesion sites (*n*). This formulation is important, because it allows peripheral adhesions to stochastically initiate ‘hot spots’ of signaling, which then dissipate if the local adhesions are not maintained. Consistent with the **p** balance, generation is saturable with respect to *a*, according to the constant parameter, *β*_*a*_. The concentration of the inactive/precursor form, *b*, is assumed to be spatially constant and determined by mass conservation, with constant parameter *b*_0_ = 1 set as the natural concentration scale.

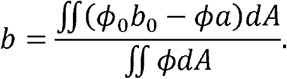

This assumption is reasonable if, for example, the inactive/precursor form of the critical signaling molecule were approximately uniform throughout the cell.

### Model modifications

Relative to the base model presented above, two modifications are presented. One is to account for the removal of ECM ligand from the surface by the migrating cells. This was incorporated in the model as a simple temporal decay of the fractional occupancy of the surface starting at *t* = *t*_*start*_.

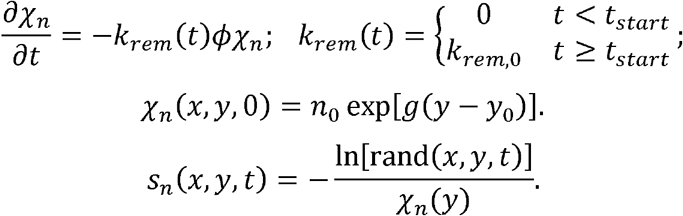

After the previously occupied sites are chosen for relegation, the value of *N*_*n*_(*t*) is updated to determine the (decreasing) number of previously unoccupied sites to switch over.

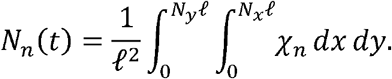

The other modification explored here is the addition of an orthogonal, chemotactic gradient in the *x*-direction. This was implemented simply by the following replacement in the generation term of the activator (*a*) balance.

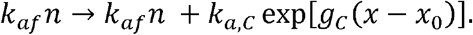

The contribution of the chemotactic gradient is determined by the magnitude and relative steepness parameters, *k*_*a,c*_ and *g*_*c*_, respectively; *x*_0_ is the *x*-coordinate of the initial cell centroid.

### Model parameterization

Dimensionless haptotactic gradient (input) parameters – the reference adhesion-site fraction, *n*_0_; and the gradient steepness, *g*, with spatial unit set to the initial radius of the cell (subsequently scaled to 20 µm to match a typical dimension of a fibroblast, spread on an adhesive surface) – were varied and expressed as percentages in the figures. In the phase field model, most parameters were fixed, based on values found suitable in published phase field models (39–41, 64) and to give suitable time scales of the reactions relative to transport and cell migration speed: Γ_*ϕ*_ = 0.5, *ϵ* = 0.05, *σ*0 1, and *α* 1; *D*_*p*_ = 3, *β* = *β* = 0.1, *k*_*pr*_ = *k*_*ar*_ = 3, and *k*_*pr*2_ = *k*_*ar*2_ = 10. The value of the dimensionless activator diffusivity, *D*_*a*_ = 1, is based on experiments (59, 65). This left three adjustable parameters (*k*_*af*_, *k*_*pf*_, and *µ*), which were varied together to identify a set of values yielding suitable signaling feedback and cell shape dynamics and qualitatively realistic cell migration tracks, while maintaining continuity of the phase field during simulations: *k*_*af*_ = 100, *k*_*pf*_ = 30, and *µ* = 1. Regarding the model modifications, low and high removal rates correspond to *k*_*rem*,0_ = 0.03 and 0.1, respectively; in simulations with a chemotactic input, *k*_*a,c*_, = 30, and *g*_*c*_ = 0, 0.05, or 0.1 as indicated in the figures.

### Python implementation – stochastic adhesion network

To maintain common sets of random numbers, to be used as replicates for all conditions, multiple random number matrices, of size number of voxels by number of time steps, were populated by “numpy.random.randoms” function, each random number determining the aforementioned ranking value, *s*_*n*_. To randomly turn off the on voxels after each timestep Δ*t*, a second random permutation matrix of size *N*_*n*_ by number of timesteps was assigned and saved. The matrix, *flip*, contains columns of 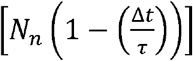 ones and (Δ*t*/*τ*)*N*_*n*_ zeros expressed in a random order using “numpy.random.permutation” function. A column is used at its respective time to turn off a random (Δ*t*/*τ*)*N*_*n*_ adhesions from the set of on voxels. When an adhesion is turned over, the voxel is reassigned a random number (and *s*_*n*_) for the next time iteration.

For the simulations with ECM removal, two additional random permutation matrices were generated using numpy.random.permutation to store the shuffled voxel index (row major index of a particular voxel) of ones and zeros, respectively, in each column of the *flip* matrix. The matrices, *ones* and *zeros*, assist in cases where the total adhesions *N*_*n*_ and turnover (Δ*t*/*τ*)*N*_*n*_ are updated based on the decreasing fractional occupancy *χ*_*n*_. The logical operation that governs this is as follows.

1. If the value of *N*_*n*_ has decreased by 1 from the previous timepoint, one more adhesion site is turned off.
2. If *N*_*n*_(*t*) < *N*_*n*_ (0),the column of *flip* must be shortened by removing *N*_*n*_ (0) − *N*_*n*_ (*t*) ones from the column.
3. If the rounded value of (Δ*t*/*τ*)*N*_*n*_ has decreased by 1, one fewer adhesion must turn off at the current timepoint.
4. If (Δ*t*/*τ*)*N*_*n*_(*t*) < (Δ*t*/*τ*)*N*_*n*_ (0), the column of *flip* must be shortened by removing (Δ*t*/*τ*) (*N*_*n*_ (0) − *N*_*n*_ (*t*) zeros from the column.

The random number matrices are read into memory and accessed by the main code to update the values of the score.

### Python implementation – phase field initialization

To initialize the phase field, *ϕ* = 1 is assigned within a unit circle of radius 1, centered at coordinates (*x*_*i*_,*y*_*i*_), followed by relaxation without the protrusion term for 50 time steps to achieve the characteristic profile of the transition region (35, 66). This initial condition is fed into the system of differential equations. To initialize the activator equation, the initialized phase field is multiplied by the constant, *k*_0_/(*k*_*ar*_*b*) The protrusion vector is initially zero. The phase field equations are solved on an 8×8 unit box with 0.05×0.05 voxel size. The 8×8 box is inset inside the 20×20 larger domain, in which the adhesion occupancy is modulated on a 0.1×0.1 voxel size. In the simulations without a chemotactic gradient, varied (*x*_*i*_,*y*_*i*_) were (6,8), (10,8), and (14,8), yielding independent replicates using the same random number set. In the simulations with both chemotactic and haptotactic gradients, (*x*_*i*_,*y*_*i*_) were fixed at (10,8) because of the *x*-dependence of the chemotactic input.

### Python implementation – solution procedure

The model was solved in a 4-step process, with a FOR loop to iterate through 20,000, 1 s (Δ*t*/2) timesteps. The relevant variables (*ϕ, a, p*_*x*_, *p*_*y*_, *χn*)and the adhesion matrix) are stored in a cell array to be passed to the four function files. First, the differential equations governing *ϕ, a, p*_*x*_, *p*_*y*_ are solved using fipy eq.solver with the LinearPCGSolver module and a tolerance of 1E-10. The solver is allowed to iterate 10 times before splitting the timestep in half and solving twice; this only occurred for the first few timesteps in cases of high gradient and high adhesion density. Second, the results are passed back to the main FOR loop to update the fractional occupancy, *χ_n_*; this step is only relevant for simulations with ECM removal. Third, if the loop is on an even iteration (every 2 s) another function file is triggered to update the adhesion matrix for the next column of the *flip* vector, as described above. Lastly, the smaller 8×8 box must be periodically moved relative to the 20×20 box. To do this, the centroid position of the phase field in the 8×8 box is checked relative to the center of the box. If the cell centroid has moved greater than 0.5 in *x* or *y*, the final function is triggered to recenter the phase field centroid and update the adhesion matrix accordingly.

### Cell migration metrics

For each simulated cell, the position of the cell centroid may be sampled at a chosen ‘microscopy’ interval *M* Δ*t*, and we index the corresponding time intervals as *i* = 1,2, …, *m*. The vector of centroid movement from image *i* to image *i* +1 is defined as **c**_*i*_. From these data, one can calculate the following quantities.

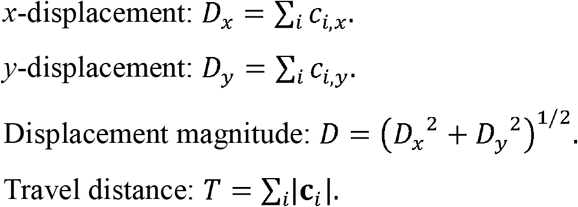

The classic metric of tactic fidelity, the Forward Migration Index (FMI), with values [−1,1], is expressed as the ratio of displacement in the direction of the gradient to the travel distance. For haptotaxis in the *y*-direction, and (in some cases) chemotaxis in the *x*-direction, we define the Haptotactic Index, *HI* = *D*_*y*_/*T*, and the Chemotactic Index, *CI* = *D*_*x*_/*T*. Another classic metric, applicable to cell migration more generally, is the Persistence ratio, defined as *D*/*T*, with values [0,11]. Based on these definitions, the different versions of the FMI may be composed as the product of the Persistence and either the sine or cosine of the Cartesian angle *θ* of the displacement vector.

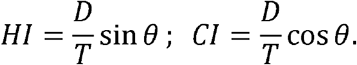

Noteworthy here is the model variation in which cells remove ECM protein from the surface and thus alter the haptotactic gradient as they migrate (by degrading the probability of adhesion site formation where the cell has resided). Experimentally, if one were able to quantify the density of ECM over time, the dynamic changes of the ECM gradient landscape could be accounted for in the tactic fidelity; however, for the semantic context of this analysis, we define the haptotactic cue here as the initial gradient.

### Experimental – cell culture

Stable expression of Paxillin-EGFP was established in the IA32 fibroblast line (67) by retroviral transduction as described (68, 69). The Paxillin-EGFP coding sequence was obtained from Addgene plasmid #15233 (70) and cloned into the pBM-IRES-Puro vector. The transduced cells were transfected for transient expression of mScarlet-AktPH using jetOPTIMUS transfection reagent (Sartorius), according to the manufacturer’s protocol. The mScarlet-AktPH C1 vector was generated by Gibson assembly and PCR to exchange EGFP from the EGFP-AktPH C1 vector described previously (57). Cells were cultured at 5% CO_2_ and 37°C in DMEM supplemented with 10% FBS and 1% penicillin-streptomycin-glutamine (PSG). For microscopy, cells were seeded at a density of 5000 cells in 1 mL of Fluorobrite DMEM supplemented with 3% FBS and 1% PSG and allowed to spread for 30 min prior to imaging. All cell culture reagents were acquired from Thermo Fisher Scientific. The cells were seeded on a dish coated with 600 µL solution of 1 µg/mL human plasma fibronectin (Fisher Scientific, #CB40008), diluted from a high concentration stock in PBS, spread over the glass bottom of a Mattek 20 mm glass bottom dish for 15 min. Cells were maintained on the scope by using an incubation system made from a house-built plexiglass box with ports for tubing connecting the AIRTHERM-SMT heater with humidifier control. The temperature was maintained at 36 °C and relative humidity at 40%. The CO_2_ was maintained at 5% using a tuned BioSpherix ProCO2 P120 Compact CO2 Controller. To minimize evaporation, 2 mL of mineral oil was added to the dish after submerging the objective.

### Experimental – TIRF microscopy

Live-cell imaging was performed using a prism-based TIRF microscope built on an Axioskop 2 FS stand (Ziess) outfitted with an x-y stage and CRISP autofocus system (Applied Scientific Instrumentation). TIRF illumination was achieved using a 60 mW, 488-nm sapphire laser (Coherent) and a 100 mW, 561-nm Cobolt 06-01 series DPL laser (Hübner Photonics). Exposure time for each was 1 s, and images were acquired every 10 s using an ORCA ER cooled CCD camera (Hamamatsu) fitted to the scope with a 0.63X camera mount and Metamorph software (Molecular Devices). A 20X water immersion objective (Zeiss) was used.

## ACKNOWLEDGMENTS

We acknowledge Shawn Van Bruggen for constructing the pBM-Paxillin-EGFP-IRES-puro plasmid. Research reported in this publication was supported by the National Institute of General Medical Sciences under award no. R01 GM141691. The content is solely the responsibility of the authors and does not necessarily represent the official views of the National Institutes of Health.

